# SPECIES DISTRIBUTION PROJECTIONS UNDER INTERNAL CLIMATE VARIABILITY REVEAL MULTIPLE PLAUSIBLE FUTURES REQUIRING FLEXIBLE CLIMATE-READY DECISIONS

**DOI:** 10.64898/2026.03.11.711202

**Authors:** Iván Felipe Benavides Martínez, Rachindra Mawalagedara, Kartik Aggarwal, Arnob Ray, Andrew J. Allyn, Katherine E. Mills, Auroop R. Ganguly

## Abstract

Traditional biodiversity projections using species distribution models (SDMs) assume that a given emissions pathway implies a largely predictable ecological response, yet how internal climate variability (ICV) translates into SDM outcomes remains largely uncharacterized. By projecting distributions for 34 marine and terrestrial species across 100 initial-condition members from CESM2-LENS2, we show that ICV alone produces qualitatively different outcomes, including reversals in projected range shifts. We classify these outcomes into four decision-relevant cases of increasing ICV susceptibility and find a pronounced marine-terrestrial divide that persists across time horizons. Responsiveness to ICV is not consistently explained by traits or local climate variability amplitude, implying it is atleast partly ecological in nature. Robust biodiversity planning therefore requires decisions stress-tested across distributions of plausible climate realizations rather than single best-estimate maps.

## 1 Introduction

Species distribution models (SDMs) are widely used to project climate-driven range shifts and guide climate-ready management (Robinson et al., 2017; Franklin, 2023; Benavides et al., 2024). But when SDMs produce a single best-estimate projection, what confidence is warranted where the driving climate trajectory is only one of many plausible realizations? Such single projections result from forcing SDMs with either a single Earth System Model (ESM) simulation or ensemble-mean climate fields averaged across realizations differing only in their initial conditions (ICs) (Kay et al., 2015; Maher et al., 2021), both of which collapse internal climate variability (ICV), the divergence among ESM realizations arising from sensitivity to initial conditions, into a single trajectory (Freer et al., 2018). The key unresolved issue is not whether ICV exists (Deser et al., 2012, 2020; Deser, 2020; Mawalagedara et al., 2025), but how to systematically translate trajectory divergence into calibrated ecological inference (Beaumont et al., 2007; Jenouvrier et al., 2025; Palacios-Abrantes et al., 2022; Wilson et al., 2024). Because SDMs are nonlinear mappings from climate histories to habitat projections, differences in the sequencing of anomalies and extremes can move species across thresholds, activate lags, and result in distinct response regimes even under identical forcing. Yet current practice provides little guidance for deciding when collapsing this uncertainty is benign versus when it yields overconfident or directionally inaccurate conclusions.

ICV varies with geographic region, spatial scale, time horizon, and climate variable (Deser et al., 2012, 2020; Kumar and Ganguly, 2018; Mawalagedara et al., 2025; Ray et al., 2025), yet its ecological consequences for SDM projections remain poorly characterized. It can be especially consequential for regional and local projections over near-term to midcentury horizons (Mankin et al., 2020; Upadhyay and Bhatia, 2025), where SDMs are often used in management decisions (Guisan et al., 2013). ICV also differs in magnitude between marine and terrestrial environments (Deser et al., 2020), raising the possibility that SDM projection susceptibility to ICV may vary across major ecological groups. However, it remains unclear whether SDM projections exhibit a range of outcomes correlated to local ICV magnitude in climate predictors or whether ecological nonlinearities amplify or dampen the effect of climate signals in ways that decouple SDM spread from ICV. Without a generalizable framework for anticipating when and where ICV produces divergent ecological outcomes, single best-estimate projections cannot be assessed for reliability, and the conditions under which they mislead remain unidentified.

To address this gap, we develop a fully ICV-aware species distribution modeling and analysis framework (fig. S1) that explicitly propagates ICV from a single ESM into ecological projections, one that (i) preserves realizable climate histories rather than averaging them away, (ii) isolates ICV as a standalone uncertainty source by holding emissions forcing, climate model structure, and SDM formulation constant while varying only initial conditions, and (iii) quantifies when projected outcomes are robust or variability-dominated. Applied consistently across biological systems and evaluated using multiple decision-relevant metrics, this framework establishes a generalizable basis for comparing projection divergence (defined here as the spread across ensemble members) and identifying the conditions under which SDM outputs remain informative, become uncertainty-limited, or cease to support confident inference.

We operationalize this ICV-aware framework across 34 species (12 marine and 22 terrestrial) selected for data sufficiency, prior SDM applications, and diversity of life history traits, biogeography, and ecological niches. Future climate predictors are derived from 100 initial-condition members of the CESM2-LENS2 large ensemble (Kay et al., 2015) under SSP3-7.0 for mid-century (2031–2060) and late-century (2071–2100) periods. To ensure consistency with observed baselines while preserving internal variability, all dynamic environmental predictors predictors are bias-corrected to ERA5 and CMEMS climatologies (1981–2014) separately for each IC member. For each species, we train a single SDM using observational climate data and static predictors (e.g., elevation and bathymetry), and project this fixed model across all 100 bias-corrected IC realizations so that only the climate trajectory varies among projections. For further analysis, from each realization, we derive FOUR range-based ecological metrics: Potential Area Change (PAC), Potential Area Gain (PAG) and Potential Area Lost (PAL), and Potential Suitability Ratio Change (PSRC). Given the breadth of species, metrics, and time horizons analyzed, results presented here reflect aggregate patterns across the full ensemble. An interactive Shiny application (Benavides, 2026) enabling extended exploration of species-specific projections, additional metrics, and full IC distributions is available online (https://felipeben.shinyapps.io/species-distribution-app/).

## 2 Results

### 2.1 Divergent species projections emerge from internal climate variability

To quantify how ICV propagates into ecological projections, we projected distributions for 34 marine and terrestrial species using 100 initial-condition members from CESM2-LENS2 under identical forcing and emissions scenario. The resulting projections show that ICV alone is sufficient to generate qualitatively different outcomes in species distribution (Figs. 1 and 2A-B). We illustrate this with two examples spanning contrasting taxonomic groups and regions (Fig. 1).

**Figure 1.**
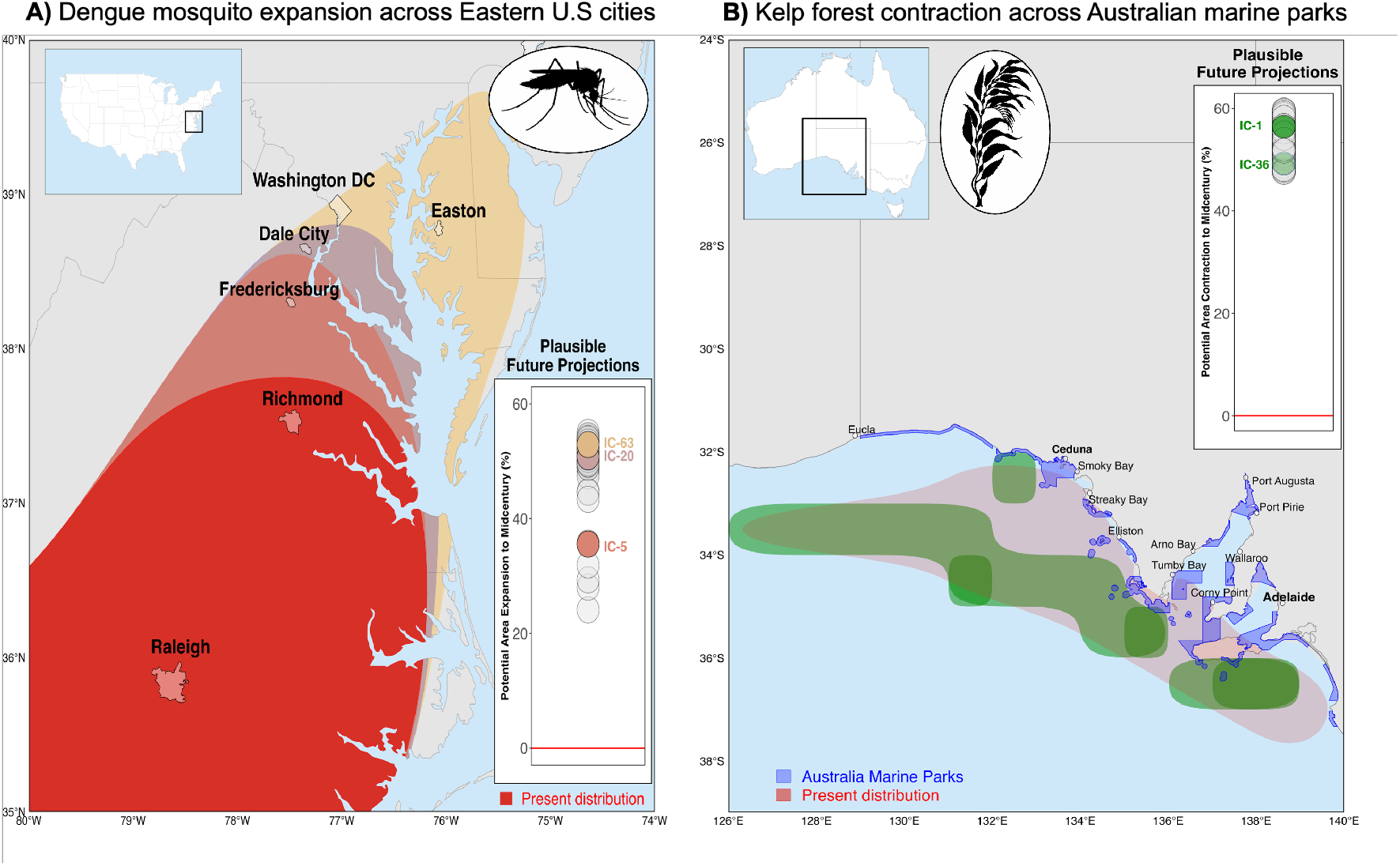
Divergent decision-relevant outcomes arising from internal climate variability. Different initial-condition (IC) realizations from the same Earth system model (CESM2-LENS2), under identical emissions forcing, generate contrasting species distribution projections with direct implications for management. (A) Mid-century projections for the dengue vector *Aedes aegypti* along the eastern United States show that alternative IC realizations produce different northward expansion limits, exposing different cities to potential transmission risk. Public-health preparedness decisions therefore may differ depending on the selected realization. (B) Late-century projections for kelp forests in southern Australia show contrasting patterns of contraction and fragmentation across IC realizations, altering which marine parks and coastal communities experience habitat loss. These examples illustrate how ICV alone can reorganize projected geographic impacts, reshaping decision landscapes for conservation, restoration, and public-health planning.

**Figure 2.**
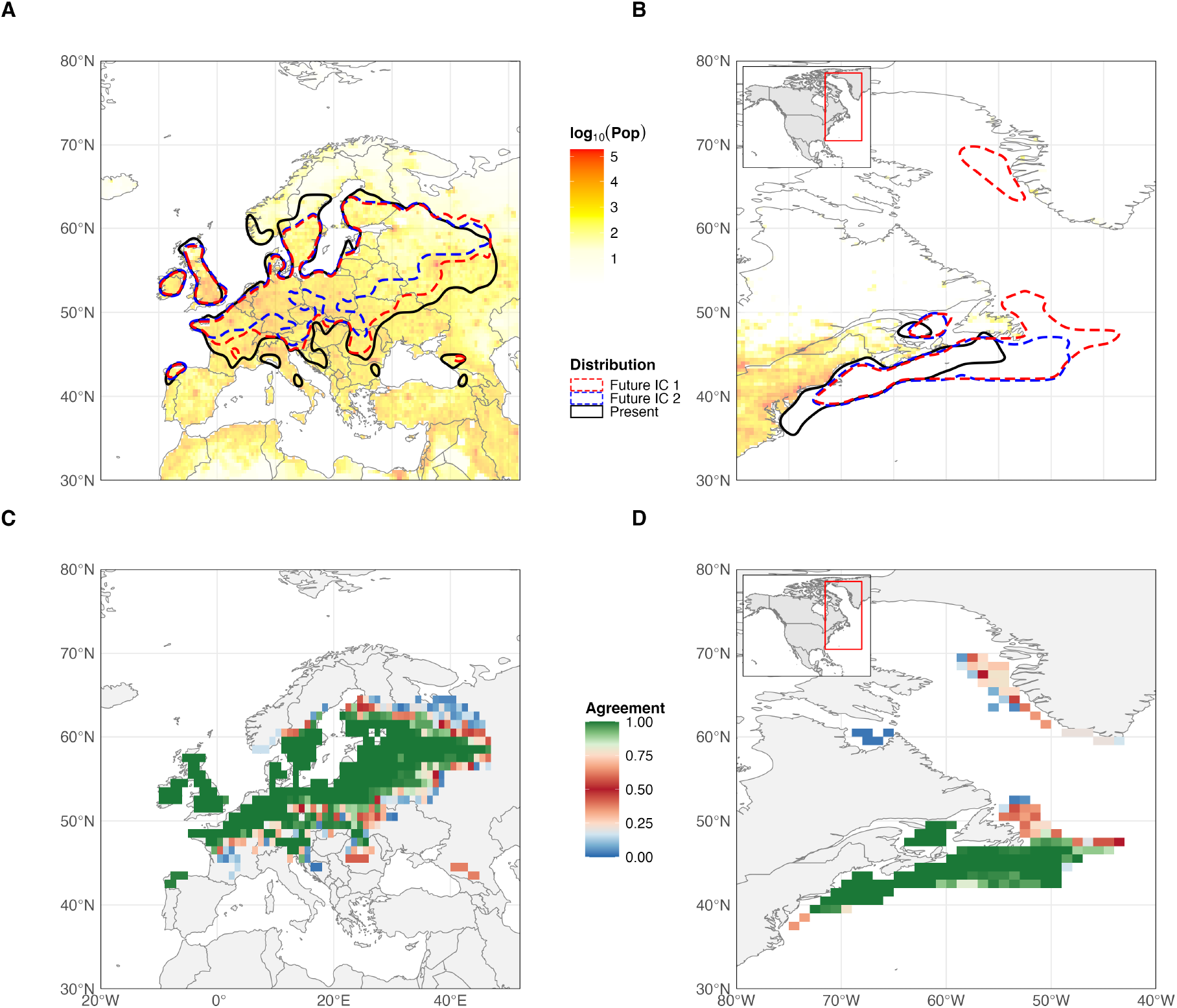
Contrasting species distribution projections and ensemble consensus under internal climate variability. Two examples from CESM2-LENS2 under SSP3-7.0 for 2071–2100 illustrate how different initial-condition (IC) realizations can produce markedly different ecological outcomes. (A-B) Present-day distributions (black outlines) are compared with two contrasting future projections from different IC realizations (red and blue), showing substantial spatial variability in projected potential range shifts for *Ixodes ricinus* (left) and *Homarus americanus* (right). Background shading shows human population density from the Gridded Population of the World dataset on a log_10_(Pop) scale. (C-D) Heat maps quantify agreement across all 100 IC realizations, where color intensity indicates the proportion of IC members projecting species presence at each location. High-agreement regions represent robust projections across ICs, whereas intermediate-consensus areas identify locations of greatest disagreement and thus strongest sensitivity to internal climate variability.

For *A. aegypti* along the eastern United States, the choice of IC determines which cities fall within potential transmission risk zones, as different realizations produce divergent northward expansion limits (Fig. 1A). In southern Australia, the choice of IC leads to substantially different degrees of contraction and fragmentation of kelp forests altering which marine parks and coastal communities face habitat loss (Fig. 1B). The choice of IC realization therefore shapes assessments of exposure, vulnerability, and conservation priority including which human populations face the greatest risk (Fig. 2A-B), with direct consequences for management and decision-making. The following results systematically quantify what these examples demonstrate qualitatively, across all 34 species, ecological metrics, and time horizons, to extract generalizable insights for more robust ecological projections (Fig. 2A-B, fig. S2A-B).

First, to quantify agreement among the 100 IC members, we calculated ensemble consensus at each grid cell as the percentage of members projecting species presence (see Methods). Fig. 2C–D illustrates spatial consensus for two species, where values near 0% or 100% indicate strong agreement among ensemble members on species absence or presence, respectively, whereas values near 50% identify locations of greatest disagreement where the ensemble is split nearly evenly. These intermediate-consensus grid cells therefore represent areas of greatest irreducible uncertainty in SDM projections. For species with relatively continuous projected ranges, lowest consensus tends to occur at range boundaries, whereas for species with highly fragmented distributions, IC disagreement is distributed throughout the range rather than concentrated at its margins.

Next, we examined IC agreement in terms of how distribution metrics (PAC, PAG, PAL, and PSRC) vary across the 100 ensemble members (Fig. 3A-B; fig. S2A-B), complementing the grid-cell-level consensus analysis. Across the ensemble, projected species distributions fall into four cases of increasing divergence, each with distinct implications for decision-making (Fig. 3, figs. S1 and S2). A given species may occupy different cases across metrics, adding a further dimension of complexity to projection interpretation.

**Figure 3.**
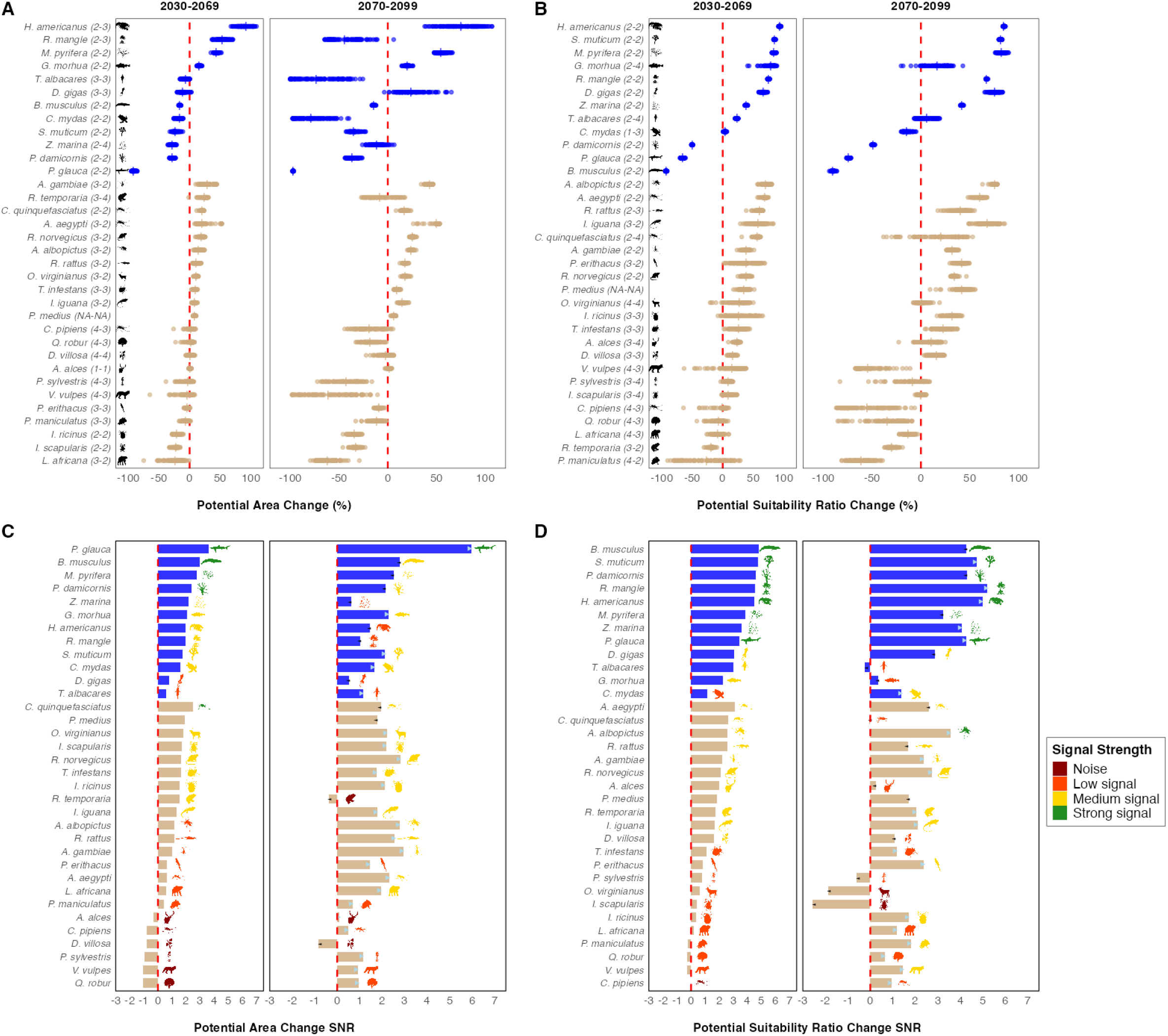
Variability in projected species distribution metrics across 100 initial-condition realizations. Species distribution model projections for 34 species across 100 CESM2-LENS2 initial-condition (IC) members under SSP3-7.0. (A) Potential area change (PAC) relative to present distributions. (B) Potential Suitability Ratio Change (PSRC). Each point represents one IC projection for a M or T species, and the numbers in parentheses indicate case classifications for mid-century (2031–2060) and late-century (2071–2100), respectively. Vertical line segments for each species indicate the conventional IC-averaged projection. (C-D) Signal-to-noise ratio (SNR) corresponding to the metrics shown in (A) and (B). Species silhouettes are colored by k-means clustering into signal categories, and arrows indicate the direction of SNR change between periods. The dashed red vertical line marks no change relative to the present distribution. Together, these results show that internal climate variability can alter both the magnitude and, for some species, the direction of projected distribution change.

We describe each case in order of increasing divergence. Case 1 encompasses species with near-neutral projected change and high agreement across ICs, indicating a robust absence of change. *A. aegypti* under PAL (fig. S2B) exemplifies this where projected area change is near neutral and tightly clustered across ICs, such that selecting any single IC yields essentially the same conclusion. Case 2 encompasses robust directional responses in which all ICs agree on both direction and magnitude of change, providing confident inference from any single realization given a sufficient ensemble size (Milinski et al., 2020). *M. pyrifera* under PAC (Fig. 3A) exemplifies this, with all 100 ICs projecting positive change within a relatively narrow range (50 to 60%). Case 3 encompasses consistent directional change but substantial divergence in magnitude across ICs, indicating agreement on the sign of response but high uncertainty in the strength of change, limiting confidence in impact and severity assessments.

*L. africana* under PAC for mid-century (Fig. 3A) illustrates this, with ICs spanning from minor decline to severe loss (down to -80%) and implying markedly different conservation responses depending on the realized climate trajectory. Case 4 encompasses directional reversals, where ICs disagree on the sign of projected change, precluding confident inference about the direction of future change itself. *V. vulpes* under mid-century PSRC (Fig. 3B) is an example for Case 4, with projections ranging from severe contraction (down to -70%) to expansion (up to 50%) of habitat suitability across ICs, rendering the ensemble mean misleading. Among these four cases, case 3 and case 4 are most consequential, as divergence across ICs either obscures the severity of change or reverses its direction entirely. Across all metrics and time periods, 44% of species classifications fall into Case 3 or case 4, indicating that nearly half of species projections are substantially altered by ICV (fig. S3). Averaging across ICs does not resolve this divergence; instead, it conceals opposing ecological outcomes. Reliance on a single IC realization or the ensemble mean can therefore reverse, cancel, or spuriously generate projected distribution change.

### 2.2 Forced signal strength partially explains but does not determine case classification

To characterize the climate signal underlying projection divergence, we computed signal-to-noise ratios (SNR) (Lehner and Deser, 2023) for each species and metric, quantifying forced habitat change relative to IC-driven spread (Fig. 3C-D and fig. S2C-D). Marine species show consistently higher SNR than terrestrial species across both metrics and periods, while terrestrial SNR varies widely and is structured by life-history traits.

Case 4 species consistently exhibit lower SNR confirming that ICV dominates where ensemble members cannot agree on direction, while Case 2 species tend toward higher SNR where the forced signal is sufficient to produce consensus on both direction and magnitude. However, within Case 2, SNR spans a wide range, indicating that consensus does not require uniformly strong forcing (fig. S4). Cases 2 and 3, which represent fundamentally different decision-making contexts, can occupy overlapping SNR ranges, reflecting that forcing strength and the structural distribution of IC projections are distinct properties. Further, Case 1 (*E*.*g*.: *A. alces*) occupies the same low-SNR space as Case 4 with low SNR values resulting from negligible change for the former and ICV dominated divergence for the latter (fig. S4).

SNR is an established diagnostic of forcing strength Deser et al. (2020), but as illustrated above, even in an application where ecological outcomes are binary, forcing strength does not capture the structural distribution of IC projections. Relying on SNR alone is therefore insufficient in this context, a limitation shared by the ensemble mean, which similarly collapses the distributional structure that determines case classification into a single statistic. Thus, our case framework operates as a decision-relevant classifier of this distributional structure, providing a complementary perspective that SNR does not access.

### 2.3 Projection divergence is structured across taxa and time horizons

Projection divergence is not randomly distributed across species. A pronounced marine-terrestrial divide emerges across all metrics and periods. Marine species overwhelmingly occupy Case 2 (80%), indicating predominantly robust and predictable responses, whereas terrestrial species distribute more evenly across cases with a substantially higher proportion occupying Cases 3 and 4 (60%) (Fig. 3A-B, fig. S2A-B).

This pattern is consistent with the known physical structure of ICV, which is systematically larger over land than ocean across the climate variables that drive SDM projections (Deser et al. (2020)), amplifying IC-driven spread in terrestrial systems relative to marine ones. SNR corroborates this mechanism: marine species consistently show higher SNR values than terrestrial species across both metrics and periods (Fig. 3C-D), confirming that the forced climate signal more reliably dominates IC-driven spread in ocean systems than on land.

Temporal dynamics further structure projection reliability and reinforce the marine-terrestrial divide. Across metrics, 34–38% of species maintain their case classification from mid-to late-century, while 50–55% change. Among those that change, transitions toward lower uncertainty occur approximately three times more frequently than toward higher uncertainty, as the forced climate signal strengthens relative to internal variability over time. This asymmetry is taxonomically structured: 75–83% of marine species retain their case classification across periods, compared with only 31–40% of terrestrial species, confirming that terrestrial projections remain more susceptible to ICV throughout the century (fig. S3).

### 2.4 Metric choice reshapes conclusions about projection divergence

Projection divergence depends not only on species and time horizon but also on the ecological metric used to quantify change. The four metrics considered capture distinct ecological processes at different spatial scales. PAC exhibits the tightest IC clustering across species, whereas PAL and particularly PSRC display substantially wider IC variability. Some species maintain consistent classifications across all metrics, representing robust responses to climate forcing. Others exhibit strong metric dependence. For example, *O. virginianus* spans multiple cases depending on the metric evaluated, while *A. aegypti* shifts case classification across metrics and time periods.

These differences do not reflect contradictions but instead highlight that area change, colonization potential, habitat loss, and suitability shifts respond differently to IC-driven variability. Species that are sessile or strongly habitat-specialized tend to show cross-metric stability (*M. pyrifera, P. damicornis, R. mangle*), whereas highly mobile generalists exhibit greater cross-metric instability (*V. vulpes, A. alces*) (Fig. 3A-B and fig. S2A-B). Projection divergence is therefore not intrinsic to species responses; it depends on how ecological change is quantified.

### 2.5 Projection divergence is not proportional to ICV magnitude

If ICV magnitude directly drives SDM disagreement, grid cells with larger climate spread would correspond to intermediate consensus where IC disagreement is the largest. However, grid cells with large or small ICV in temperature, precipitation, relative humidity, wind speed, or sea-surface temperature do not systematically align with particular consensus levels for either marine or terrestrial species (figs. S5 and S6). ICV ranges overlap substantially among consensus bins, and no consistent latitudinal structure is detected. This conclusion holds even when ICV is evaluated in multivariate climate space. Large climate spread does not, by itself, identify where ensemble members diverge in their SDM projections.

The divergence of species distribution projections is therefore not explained by the local amplitude of climate variability. Having ruled out a simple magnitude-based explanation, we tested whether how ICs are arranged within the local environmental envelope (described by the environmental predictors used in the study) explain their divergence. We quantified the separation between ICs projecting presence and absence using raw energy distance (ED) (Rizzo and Székely, 2016) in multivariate environmental space, focusing on the most influential dynamic predictors for each species (Fig. 4A–B). Across both terrestrial and marine taxa, ED is non-zero at most grid locations, indicating systematic separation between presence- and absence-projecting ICs.

**Figure 4.**
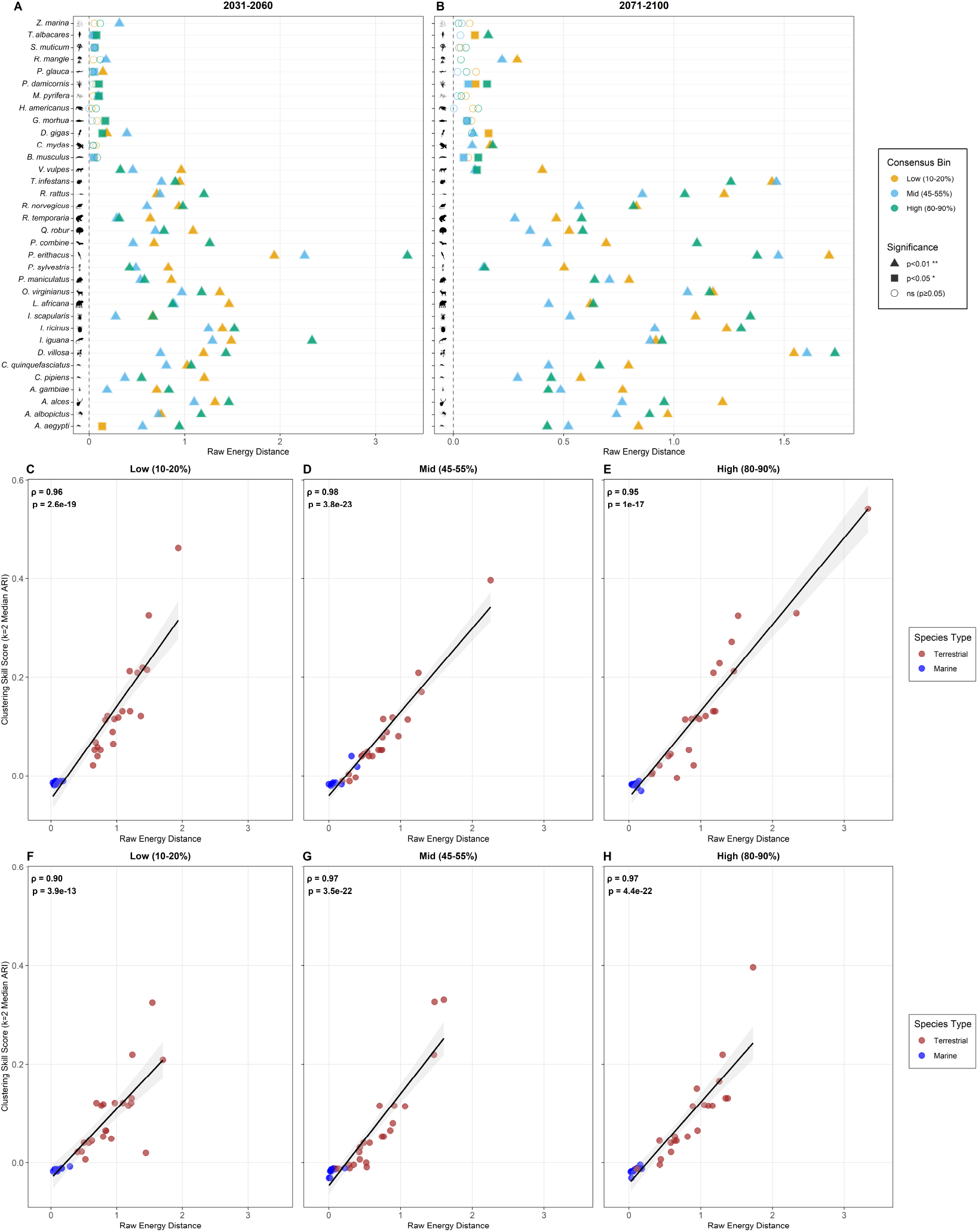
Environmental separation between presence- and absence-projecting ensemble members and its recoverability by clustering. Raw energy distance (ED) quantifies separation between initial-condition (IC) members projecting presence versus absence in the multivariate space of the four most important dynamic predictors affected by internal climate variability for each species. (A-B) ED for low (10-20%), mid (45-55%), and high (80-90%) presence-consensus bins for 2031–2060 (A) and 2071–2100 (B). Each point represents a species–bin combination, with species ordered by realm. Point colors denote consensus bins and point shapes indicate significance relative to a permutation-based null model. (C-E) Relationship between ED and clustering skill score for 2031–2060 across low, mid, and high consensus bins, respectively. (F-H) Same as (C-E) but for 2071–2100. Clustering skill is measured as the median adjusted Rand index from *k* = 2 hierarchical clustering applied to the same environmental predictors used in the ED analysis. Points correspond to species and are colored by realm. Larger ED values are generally associated with higher clustering skill, indicating that stronger environmental separation makes presence–absence groups more recoverable from environmental predictors alone, although environmental separation remains insufficient to fully reproduce SDM outcomes.

For terrestrial species, separation is consistently significant (*p <* 0.01 for all terrestrial species–bin combinations except one, which was significant at *p <* 0.05); marine species show a mixture of significant and non-significant separations, generally with smaller magnitude. Across species, ED patterns varied across low or mid or high bins, with no consistent ordering that would map consensus level to a single form of predictor-space separation. Importantly, the within-cell ICV ranges for these predictors are bounded and finite (figs. S5 and S6), yet ED shows that IC members projecting presence versus absence are often separated within this envelope. This is consistent with the possibility that nonlinear SDM responses can translate within-ensemble climate differences into divergent binary outcomes for habitat distributions.

### 2.6 Environmental predictors only partially explain distribution divergence

To evaluate whether environmental differentiation alone can recover SDM-projected presence and absence groups, we applied unsupervised hierarchical clustering to the same environmental predictors and compared resulting clusters with SDM classifications. Clustering skill correlates strongly with ED (*ρ ≈* 0.90-0.98, *p ≪* 0.001; Fig. 4C–H), confirming that environmental separation is real and structured. However, even in the best cases, clustering of raw environmental predictors recovers less than 60% of SDM presence or absence groupings, as measured by the adjusted Rand index (ARI) (Hubert and Arabie, 1985) indicating that clustering on raw predictors does not fully reproduce SDM outcomes.

This suggests that small IC-driven differences in environmental predictors might be amplified through nonlinear SDM response surfaces, including interactions among dynamic variables and static constraints such as elevation and ecoregion. Ecological divergence exceeds what would be expected from environmental separation alone. It is possible that while ICV influences environmental space, SDMs nonlinearly amplify these interplays into divergent species projections.

## 3 Discussion

Conventional SDM practices collapse ICV into a single trajectory, implicitly treating ICV as noise to be averaged away. However this approach obscures irreducible uncertainty in climate projections due to ICV that persists regardless of the choice of emissions scenario and ESM (Mawalagedara et al., 2025) and that is not eliminated by downscaling to finer spatial scales (Xie et al., 2015). Our results show instead that the inclusion of ICV expands the decision space beyond a “best guess” future towards an envelope of multiple plausible futures that could be qualitatively distinct that cannot be recovered from any single realization or ensemble mean. This expansion is most pronounced at mid-century, the time horizon most relevant for near-term management decisions but persists into end-century for a substantial subset of terrestrial species, confirming that the expanded decision space is not a transient feature of near-term projections alone. This expansion and its persistence are not a barrier to action but an indication of the need for more flexible thinking and actions. Our case framework, which can be extended beyond SDMs to any application with binarized outcomes (e.g. heatwave or coldsnap occurrences), positions decision-makers within this expanded space by translating ensemble behavior into four actionable pathways. These include targeted adaptation where direction and magnitude are robust (Case 2), flexible policy design where direction is clear but magnitude is not (Case 3), and no-regrets measures where direction itself is unresolved (Case 4) (Fig. 5). However, its important to note that, at very high spatial resolutions, even cases 1 and 2 which offer the most tractable decision contexts overall may exhibit locally meaningful divergence. Therefore, based on need, climate ready actions that are robust across multiple plausible futures requires stress-testing decisions across the full envelope of plausible futures rather than optimizing for a single trajectory, an approach consistent with robust decision-making frameworks developed for deep uncertainty (Lempert, 2003; Lempert and Groves, 2010).

**Figure 5.**
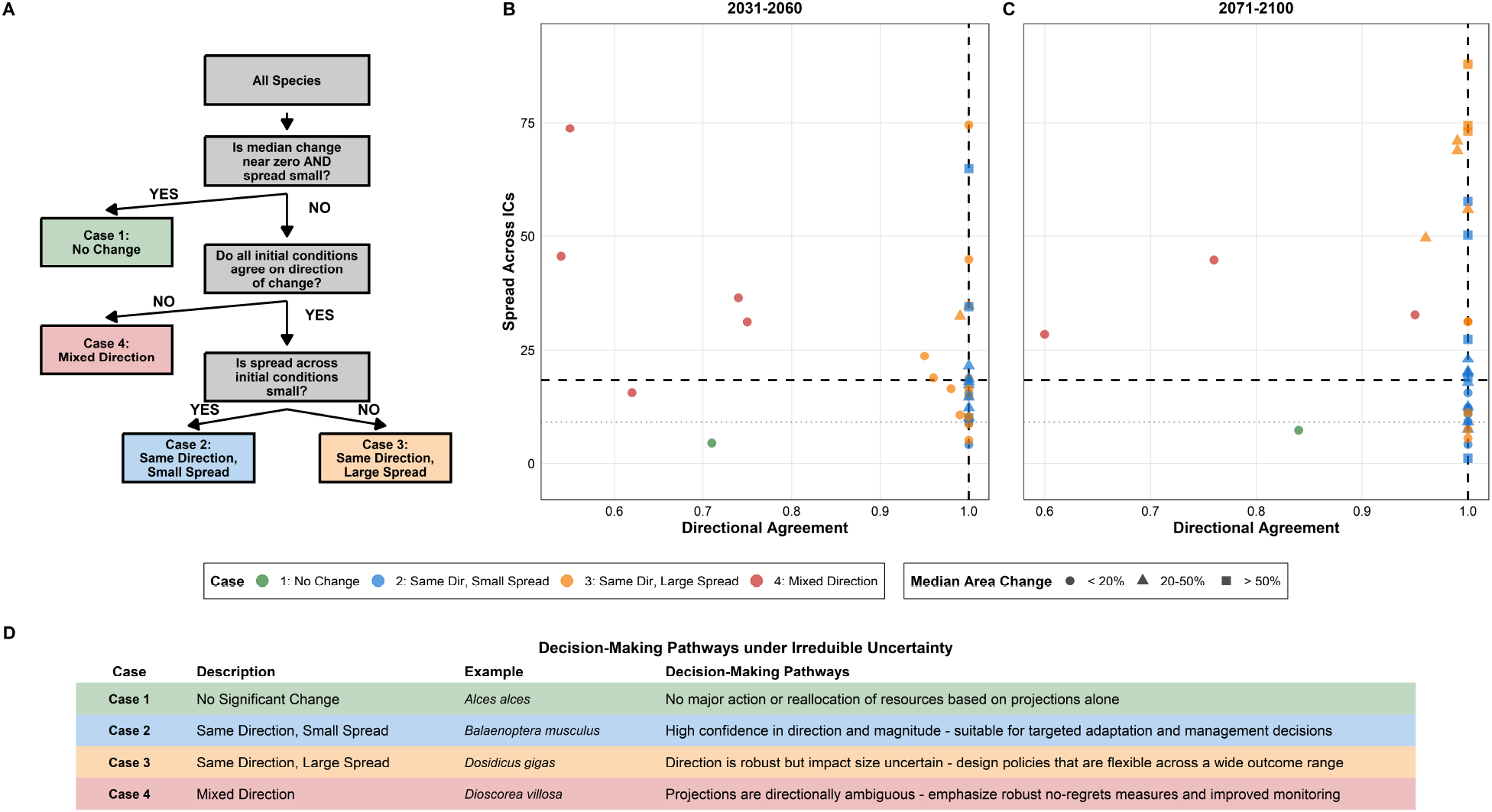
Decision-making framework for species distribution models under irreducible uncertainty. (A) Conceptual decision tree sequentially classifying outcomes from ensemble projections based on agreement in predicted area change across initial conditions. Species are evaluated first for negligible change, then for directional consensus, and finally for magnitude uncertainty. (B-C) Decision space for mid-century (2031–2060) and late-century (2071–2100) projections, showing 34 species positioned by directional agreement (x-axis) and spread across ensemble members (y-axis). Point color indicates classification case (Cases 1–4), separated by corresponding threshold lines. Point shape indicates magnitude of median change. (D) Decision-making pathways linking projection characteristics to management strategies. Case 1 species show negligible projected change and support baseline monitoring. Case 2 species exhibit high-confidence directional and magnitude signals and support targeted interventions. Case 3 species show robust directional trends but uncertain magnitudes, requiring flexible policy design. Case 4 species display directional ambiguity, warranting no-regrets approaches and enhanced monitoring.

## 4 Materials and Methods

### 4.1 Species occurrence data

We analyzed 34 species spanning marine (n = 12) and terrestrial (n = 22) realms, selected to represent diverse taxonomic groups, spatial extents (global and regional), and socioecological relevance, including conservation priority species, commercial fisheries, forestry resources, disease vectors and zoonotic reservoirs, and invasive species of biosecurity concern (table S1). Species were required to meet four criteria: (i) sufficient occurrence records to support robust species distribution models (SDMs), (ii) documented prior use in SDM studies, (iii) relevance to real-world decision-making, and (iv) coverage of contrasting ecological niches and life-history strategies.

Occurrence records were obtained from the Global Biodiversity Information Facility (GBIF) using the R package rgbif (Chamberlain et al. (2025)). Records were filtered to remove observations with missing or invalid coordinates, zero-variance coordinates, or spatial inconsistencies with species habitat (e.g., marine species on land). To reduce spatial sampling bias and autocorrelation, occurrences were spatially thinned such that all records within a 1° × 1° grid cell were treated as a single presence (Boria et al. (2014)). This ensured consistency with the spatial resolution of climate predictors and prevented overrepresentation of heavily sampled regions.

### 4.2 Environmental predictors

Present-day climatologies were assembled from ERA5 (Hersbach et al. (2020)) reanalysis for terrestrial variables and from the Copernicus Marine Environment Monitoring Service (CMEMS) (E.U. Copernicus Marine Service Information (CMEMS) (year)) for marine variables. Annual means were computed for the baseline period 1981–2014. Terrestrial predictors included near-surface air temperature, total precipitation, relative humidity, and wind speed. Marine predictors included sea surface temperature, surface dissolved oxygen, surface pH, surface nutrients (nitrate, phosphate, silicate, iron), and surface currents (u and v components). Static predictors comprised elevation and ecoregion for terrestrial species and bathymetry for marine species. All predictors were resampled to a 1° global grid to match the native resolution of the CESM2-LENS2 climate projections. Continuous variables were interpolated using bilinear resampling, whereas categorical variables (ecoregions) were resampled using modal assignment to preserve class identity.

### 4.3 Climate projections and bias correction

Future climate projections were obtained from the CESM2-LENS2 single-model initial-condition large ensemble (Rodgers et al. (2021)), which provides 100 ensemble members under the SSP3-7.0 scenario. Future periods were defined as 2031–2060 (mid-century) and 2071–2100 (late-century). To align CESM2-LENS2 outputs with the observational baseline while preserving variability among ensemble members, we applied delta-method bias correction separately for each variable, grid cell, and initial-condition member (Maraun and Widmann, 2018).

For variables with approximately additive behavior, bias correction followed:

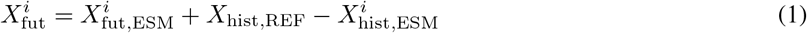

For precipitation, which is non-negative and skewed, a multiplicative delta method was used:

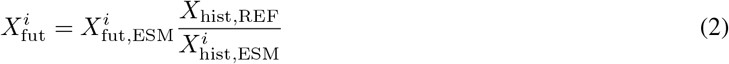

Here, 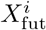 represents the bias-corrected outputs, where *i* denotes the ensemble member. *X*_hist,REF_ is the observed climatology, while 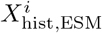 and 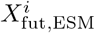 are the historical and future CESM2-LENS2 climatologies, respectively. All corrected outputs were saved as GeoTIFF or NetCDF files with filenames encoding species, variable, period, and ensemble member.

### 4.4 Species distribution modelling

Because verified absence data were unavailable, pseudoabsences were generated following Barbet-Massin et al. (2012). For each species, the number of pseudoabsences equaled the number of presences, with five independent pseudoabsence replicates. After preliminary evaluation, a surface range envelope strategy was adopted, as it maximized cross-validated performance. SDMs were fitted using XGBoost implemented in the biomod2 R package (Thuiller et al. (2025)). Models were trained using the bias-corrected climate predictors and static variables, with all predictors min–max scaled to [0,1]. Missing raster cells were filled with zero to avoid projection failures. To reduce unrealistic extrapolation, predictor stacks were cropped to a 10° buffer around the convex hull of occurrences and masked to appropriate land or ocean domains.

Models were evaluated using five-fold cross-validation and four performance metrics: True Skill Statistic (TSS), Area Under the Curve (AUC), Receiver Operating Characteristic (ROC), Accuracy, and Cohen’s Kappa. Only model runs achieving values higher or equal than 0.8 across all metrics were retained. Final ensemble SDMs were constructed by averaging retained runs for each species.

### 4.5 Projection and Ecological Metrics

Ensemble SDMs were projected to the baseline period and to each of the 100 initial-condition members for both future periods. Continuous habitat suitability outputs were exported as GeoTIFF files. Binary presence–absence maps were derived using a composite threshold defined as the mean of four cut-offs: ROC-optimized threshold, maximum sensitivity + specificity, equal sensitivity and specificity, and a quantile regression turning point. From these outputs we calculated four decision-relevant ecological metrics: Potential Area Change (PAC), Potential Area Gain (PAG), Potential Area Loss (PAL), and Potential Suitability Ratio Change (PSRC). Metrics were computed separately for each species, period, and ensemble member. To reduce sensitivity to minor fluctuations, changes within a ±10% neutrality band were excluded. For visualization, metrics were standardized to comparable ranges.

### 4.6 Signal-to-noise ratio and clustering

For each species, metric, and time period, the signal was defined as the ensemble mean across initial-condition members, and noise as the corresponding standard deviation. The signal-to-noise ratio (SNR) was computed as the mean divided by the standard deviation, with a small constant added to avoid division by zero. SNR values were standardized to range from -10 to +10, where negative values indicate variability-dominated projections. Species were classified into reliability groups using k-means clustering applied to standardized SNR values. Clustering was performed separately for mid- and late-century periods, with four clusters selected based on maximization of between-cluster variance. Clusters were labeled as: Noise, Low Signal, Moderate Signal, and Strong Signal (Fig. 3C,D and fig. S2-D).

### 4.7 Consensus, energy distance, and clustering recoverability

Species’ consensus at each grid cell was defined as the proportion of ensemble members projecting presence. Analyses were focused on cells with consensus between 10% and 90% to ensure sufficient representation of both presence and absence outcomes. To assess whether presence- and absence-projecting ensemble members occupied distinct environmental conditions, we computed raw energy distance in the multivariate space of the four most influential dynamic predictors for each species. Variables were standardized within grid cells, and balanced subsampling was applied to control for unequal group sizes.

Statistical significance was assessed using permutation tests that preserved group sizes while randomly relabeling ensemble members. To evaluate whether environmental predictors alone could recover SDM outcomes, hierarchical clustering (Ward’s D2, k = 2) was applied to the same predictor sets, and recovery skill was quantified using the adjusted Rand index across repeated balanced subsamples.

## 5 Statements and Declarations

### 5.3 Data Availability Statement

Species occurrence records used in this study were obtained from the Global Biodiversity Information Facility (GBIF) (https://www.gbif.org/). Present-day terrestrial climatologies were derived from the ERA5 reanalysis distributed through the Copernicus Climate Data Store (https://cds.climate.copernicus.eu/datasets/reanalysis-era5-single-levels-monthly-means). Future climate projections were obtained from the CESM2 Large Ensemble Community Project (LENS2) (https://www.cesm.ucar.edu/community-projects/lens2). Present-day marine environmental predictors were obtained from the Copernicus Marine Service Data Store (https://data.marine.copernicus.eu/), including global physical and biogeochemical reanalysis products. All datasets are publicly available from their respective repositories, subject to the providers’ terms of access and use.

### 5.4 Code Availability Statement

All analyses were conducted in R using the biomod2 package for species distribution modeling and ensemble forecasting (https://cran.r-project.org/package=biomod2). An interactive Shiny dashboard for exploring species-specific projections and ensemble outcomes is publicly available at https://felipeben.shinyapps.io/species-distribution-app/.

## Acknowledgments

This research was supported by AI for Climate and Sustainability (AI4CaS) of the Institute for Experiential AI (EAI), and the Alfond Foundation, both at Northeastern University, and the Gulf of Maine Research Institute. Any specific grants that need to be included??

## Supplementary Information

**Table S1.**
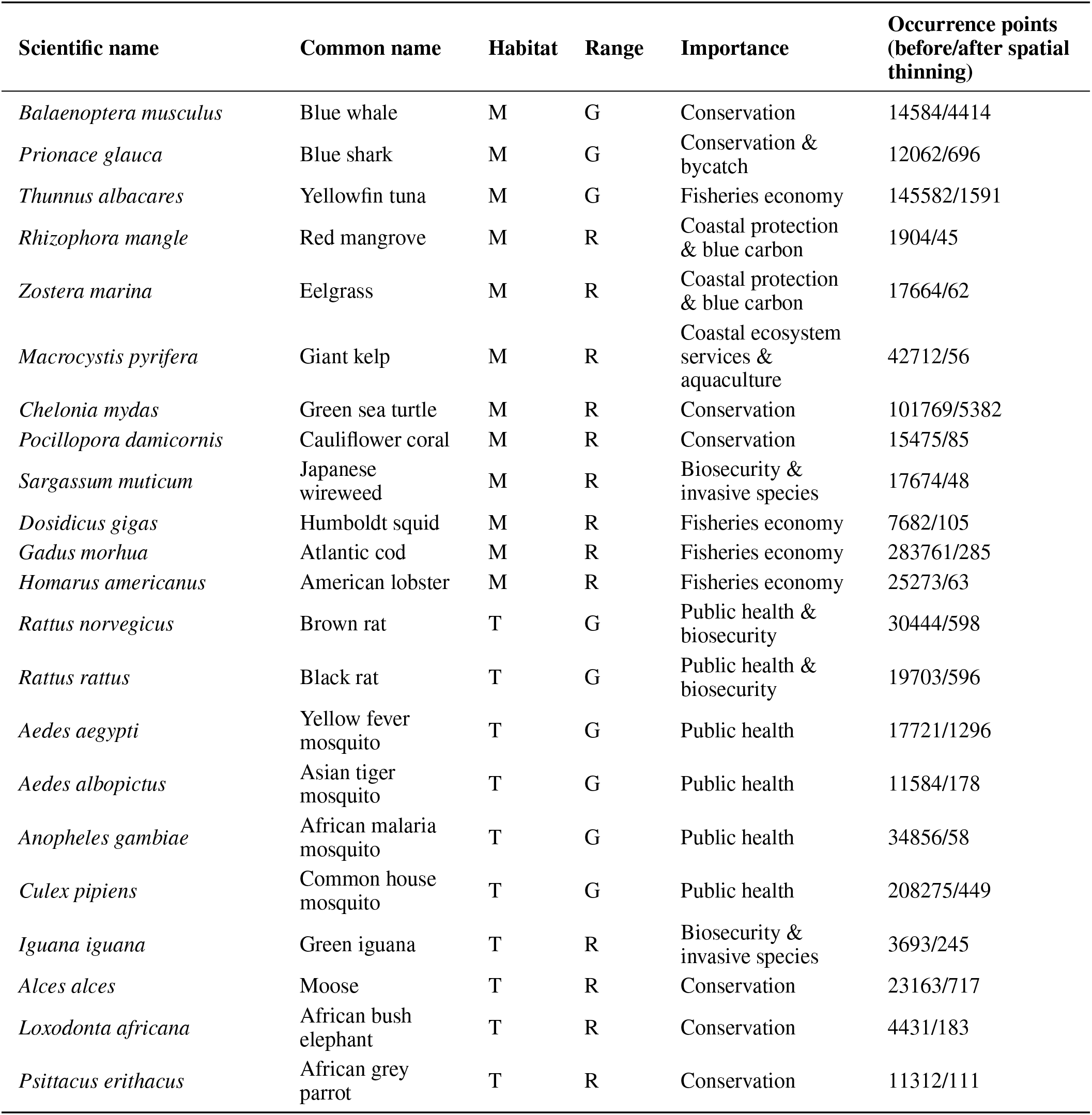
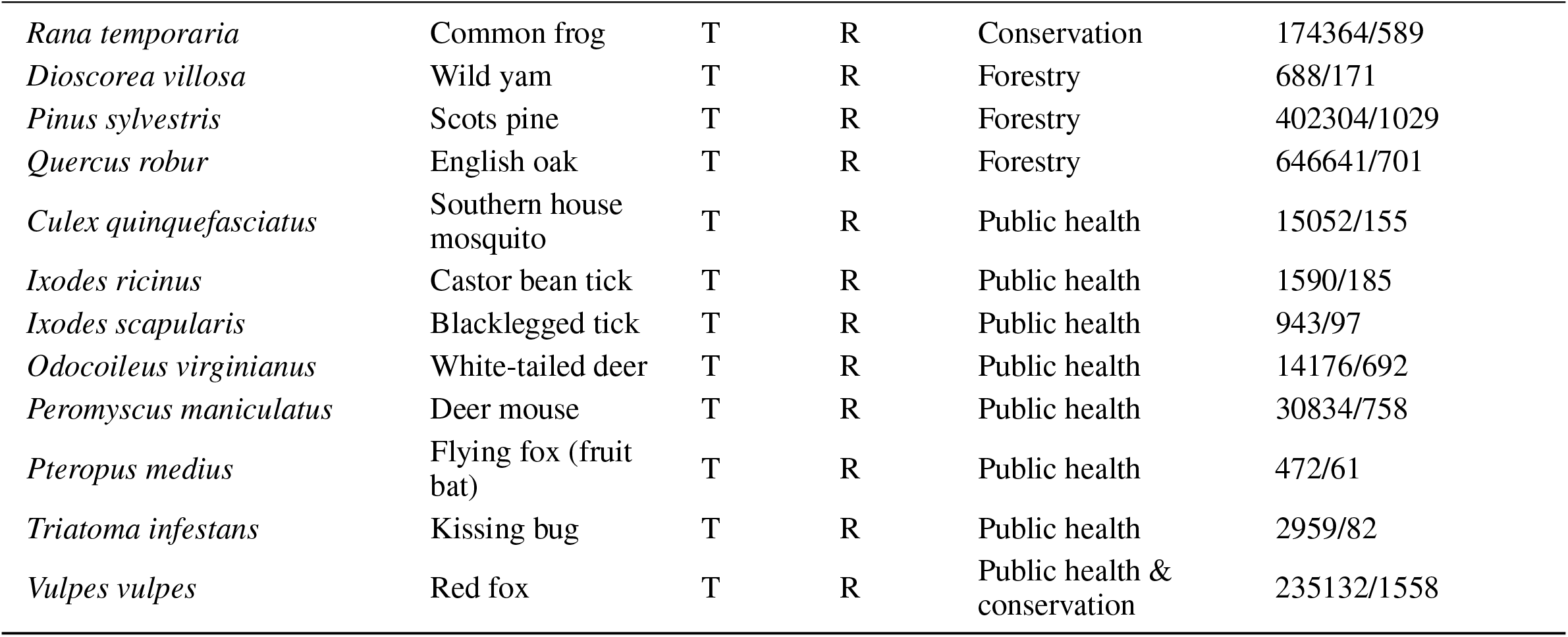
**The study includes 34 species spanning marine (M) (n=12) and terrestrial (T) (n=22) habitats, with Global (G) (n=9) and Regional (R) (n=25) distributions. Species were selected to represent diverse taxonomic groups and socioecological importance categories: conservation priority, commercial fisheries, forestry resources, disease vectors and zoonotic reservoirs, and invasive species of biosecurity concern. Occurrence records were obtained from the Global Biodiversity Information Facility (GBIF) and subjected to spatial thinning to reduce sampling bias and spatial autocorrelation while maintaining adequate sample sizes for model training. This taxonomic and functional diversity enables assessment of how initial condition uncertainty propagates across different biological systems and spatial scales.**

| Scientific name | Common name | Habitat | Range | Importance | Occurrence points (before/after spatial thinning) |
| --- | --- | --- | --- | --- | --- |
| *Balaenoptera musculus* | Blue whale | M | G | Conservation | 14584/4414 |
| *Prionace glauca* | Blue shark | M | G | Conservation & bycatch | 12062/696 |
| *Thunnus albacares* | Yellowfin tuna | M | G | Fisheries economy | 145582/1591 |
| *Rhizophora mangle* | Red mangrove | M | R | Coastal protection & blue carbon | 1904/45 |
| *Zostera marina* | Eelgrass | M | R | Coastal protection & blue carbon | 17664/62 |
| *Macrocystis pyrifera* | Giant kelp | M | R | Coastal ecosystem services & aquaculture | 42712/56 |
| *Chelonia mydas* | Green sea turtle | M | R | Conservation | 101769/5382 |
| *Pocillopora damicornis* | Cauliflower coral | M | R | Conservation | 15475/85 |
| *Sargassum muticum* | Japanese wireweed | M | R | Biosecurity & invasive species | 17674/48 |
| *Dosidicus gigas* | Humboldt squid | M | R | Fisheries economy | 7682/105 |
| *Gadus morhua* | Atlantic cod | M | R | Fisheries economy | 283761/285 |
| *Homarus americanus* | American lobster | M | R | Fisheries economy | 25273/63 |
| *Rattus norvegicus* | Brown rat | T | G | Public health & biosecurity | 30444/598 |
| *Rattus rattus* | Black rat | T | G | Public health & biosecurity | 19703/596 |
| *Aedes aegypti* | Yellow fever mosquito | T | G | Public health | 17721/1296 |
| *Aedes albopictus* | Asian tiger mosquito | T | G | Public health | 11584/178 |
| *Anopheles gambiae* | African malaria mosquito | T | G | Public health | 34856/58 |
| *Culex pipiens* | Common house mosquito | T | G | Public health | 208275/449 |
| *Iguana iguana* | Green iguana | T | R | Biosecurity & invasive species | 3693/245 |
| *Alces alces* | Moose | T | R | Conservation | 23163/717 |
| *Loxodonta africana* | African bush elephant | T | R | Conservation | 4431/183 |
| *Psittacus erithacus* | African grey parrot | T | R | Conservation | 11312/111 |

| Scientific name | Common name | Habitat | Range | Importance | Occurrence points<br>(before/after spatial<br>thinning) |
| --- | --- | --- | --- | --- | --- |
| *Rana temporaria* | Common frog | T | R | Conservation | 174364/589 |
| *Dioscorea villosa* | Wild yam | T | R | Forestry | 688/171 |
| *Pinus sylvestris* | Scots pine | T | R | Forestry | 402304/1029 |
| *Quercus robur* | English oak | T | R | Forestry | 646641/701 |
| *Culex quinquefasciatus* | Southern house<br>mosquito | T | R | Public health | 15052/155 |
| *Ixodes ricinus* | Castor bean tick | T | R | Public health | 1590/185 |
| *Ixodes scapularis* | Blacklegged tick | T | R | Public health | 943/97 |
| *Odocoileus virginianus* | White-tailed deer | T | R | Public health | 14176/692 |
| *Peromyscus maniculatus* | Deer mouse | T | R | Public health | 30834/758 |
| *Pteropus medius* | Flying fox (fruit<br>bat) | T | R | Public health | 472/61 |
| *Triatoma infestans* | Kissing bug | T | R | Public health | 2959/82 |
| *Vulpes vulpes* | Red fox | T | R | Public health &<br>conservation | 235132/1558 |

**Figure S1.**
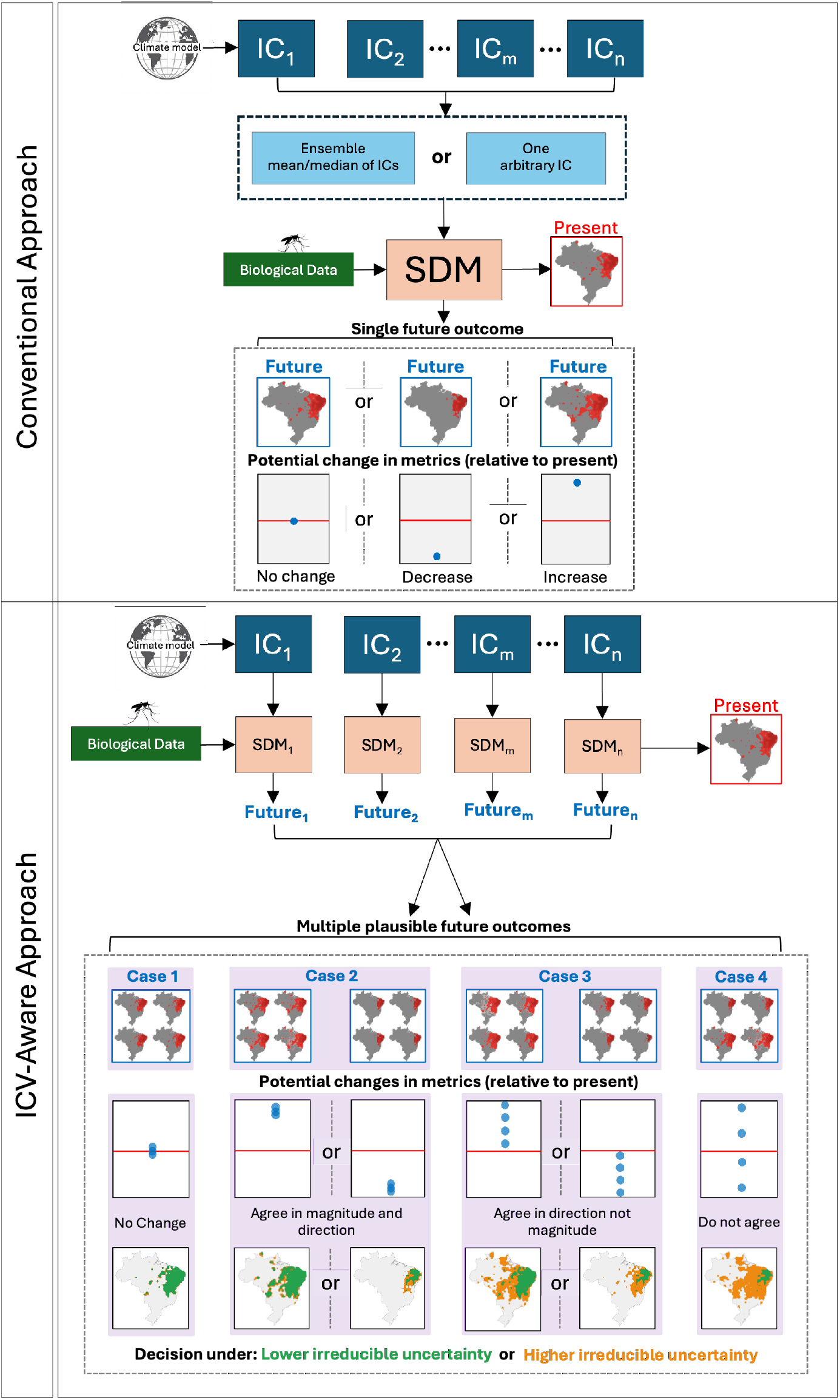
Conventional and ICV-aware species distribution modeling frameworks. In conventional SDMs, ICV is implicitly collapsed by selecting a single initial condition or by averaging across initial-condition ensemble members from a single Earth system model and emissions scenario, yielding a single deterministic future projection. In contrast, an ICV-aware approach propagates variability across initial conditions through parallel SDMs, generating multiple plausible future outcomes for the same forcing and model structure. Comparing these outcomes across ensemble members reveals whether projected changes are consistent in direction and magnitude or instead diverge, distinguishing decision contexts characterized by lower versus higher irreducible uncertainty.

**Figure S2.**
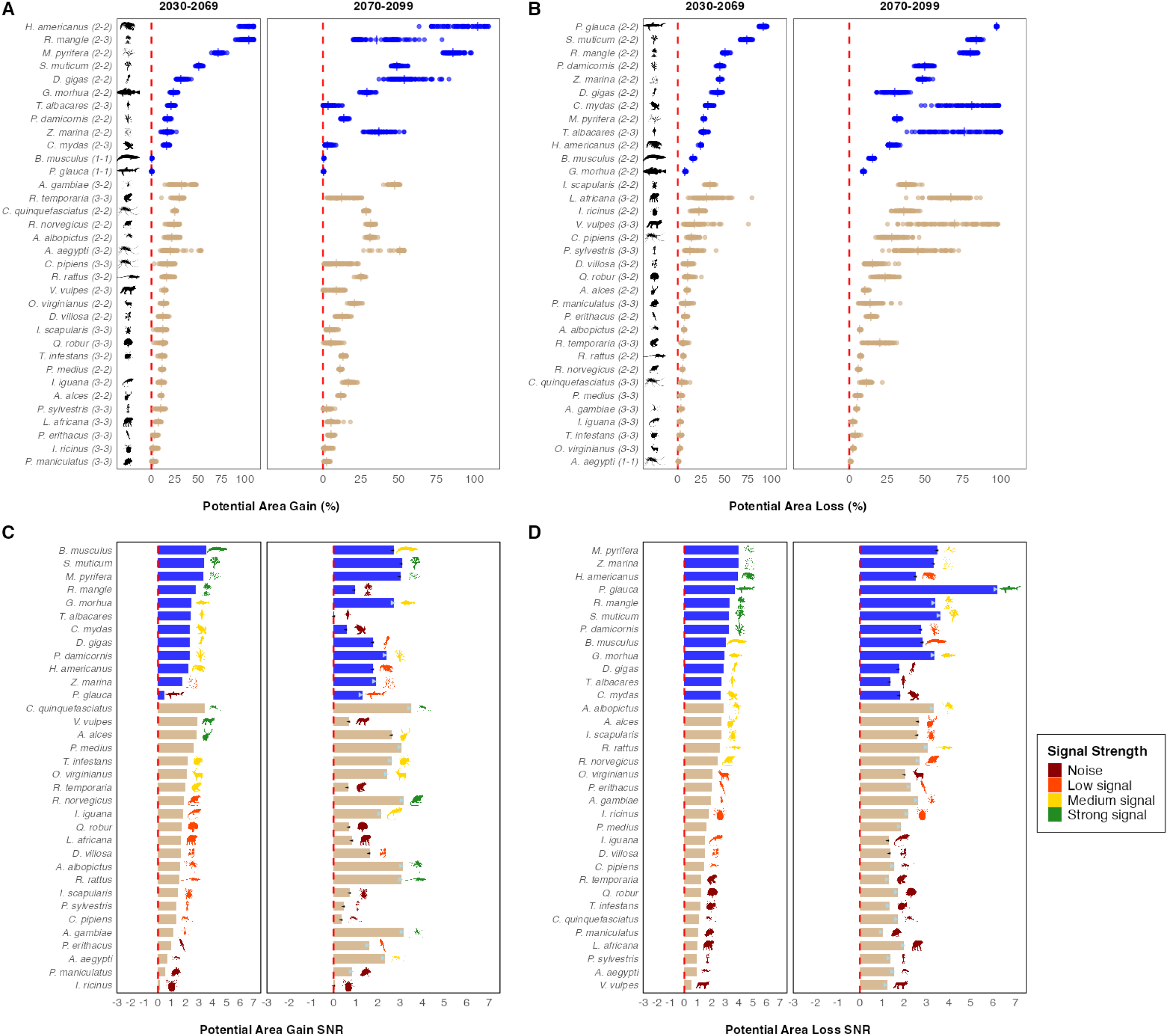
Variability in projected SDM metrics for 34 species across 100 CESM2-LENS2 initial-condition members. (A-B) Potential area gain and loss relative to present distributions. In (A-B), each point represents one IC projection for each marine and terrestrial species, while numbers in parentheses indicate case classifications for 2031–2070 and 2071–2100, respectively. Vertical line segments for each species indicate the conventional IC-averaged projection. (C-D) Signal-to-noise ratio associated with the metrics in (A) and (B); species silhouettes are colored by k-means clustering into signal categories. Arrows indicate the direction of SNR change between periods. The dashed red vertical line indicates no change compared to the present distribution.

**Figure S3.**
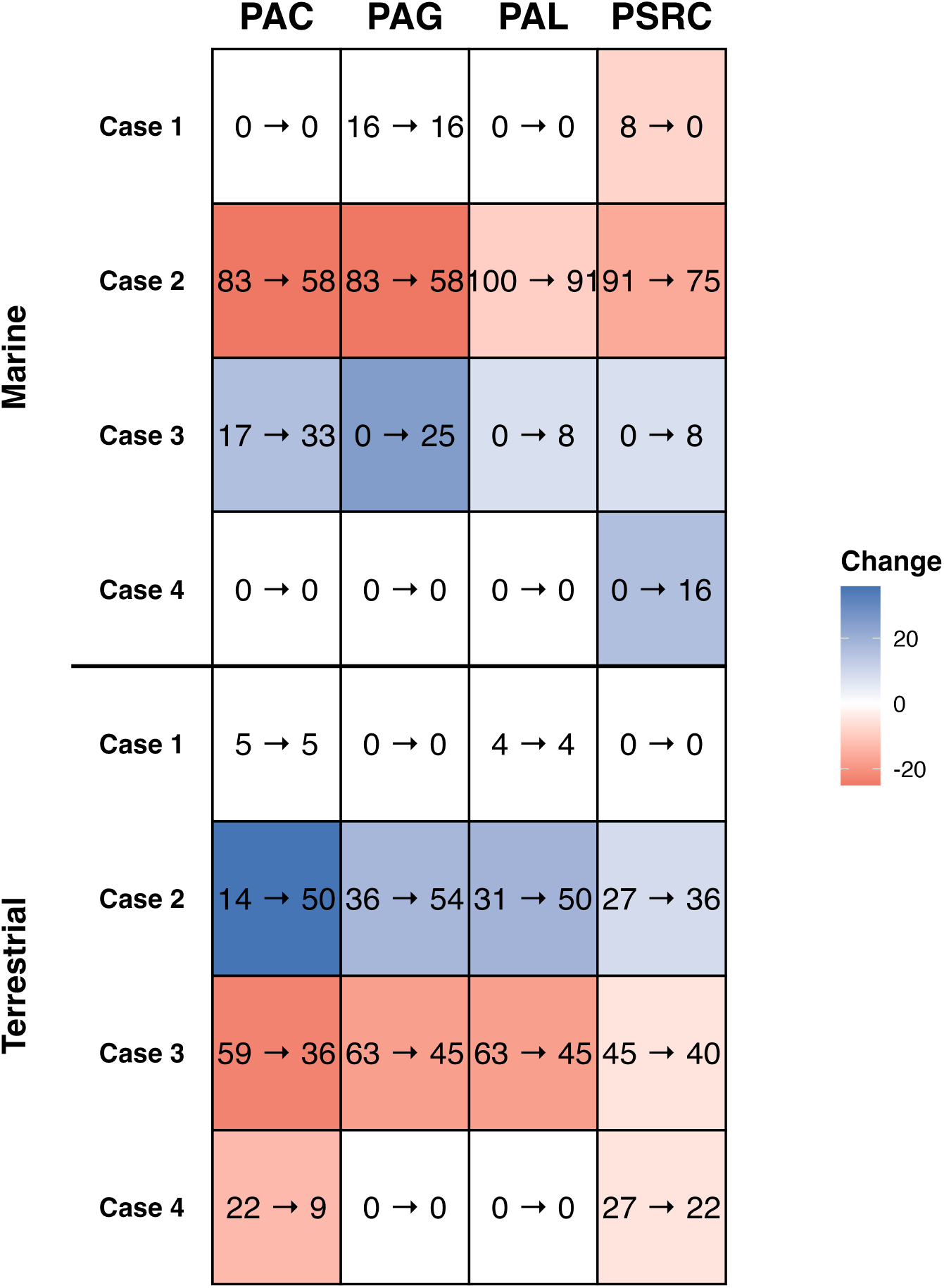
Distribution of species projections across outcome cases under internal climate variability. Percentages indicate the fraction of species classified into four projection outcome cases (Cases 1–4) for Marine and terrestrial species across ecological metrics. Cell colors indicate the direction and magnitude of change in the proportion of species from mid-to late century.

**Figure S4.**
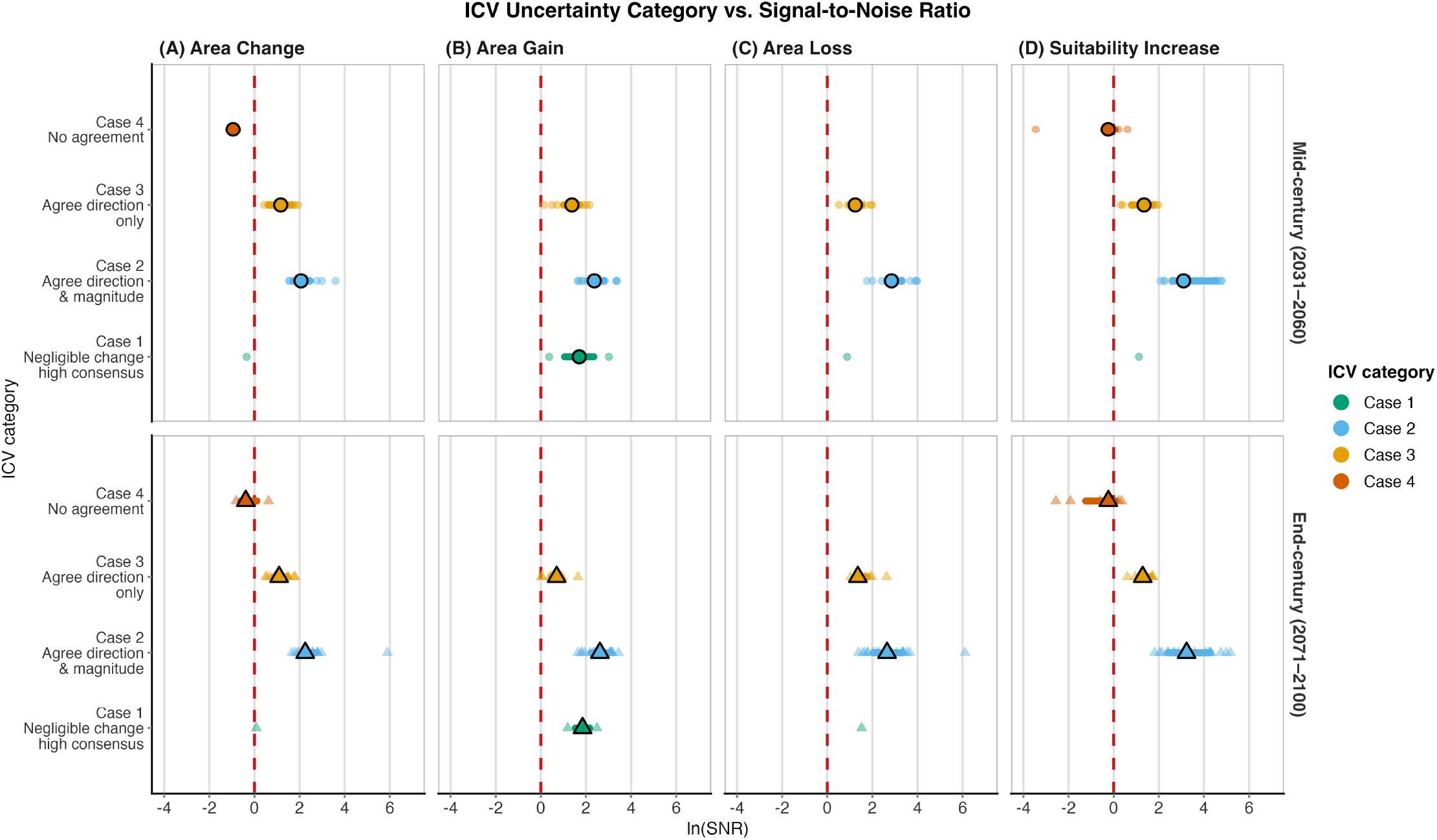
Signal-to-noise ratios associated with ICV projection categories. Species projections classified into four ICV outcome cases are plotted against the natural logarithm of the signal-to-noise ratio for multiple distribution metrics. Higher SNR values indicate projections dominated by climate-forced signal, whereas lower values indicate outcomes strongly influenced by internal climate variability.

**Figure S5.**
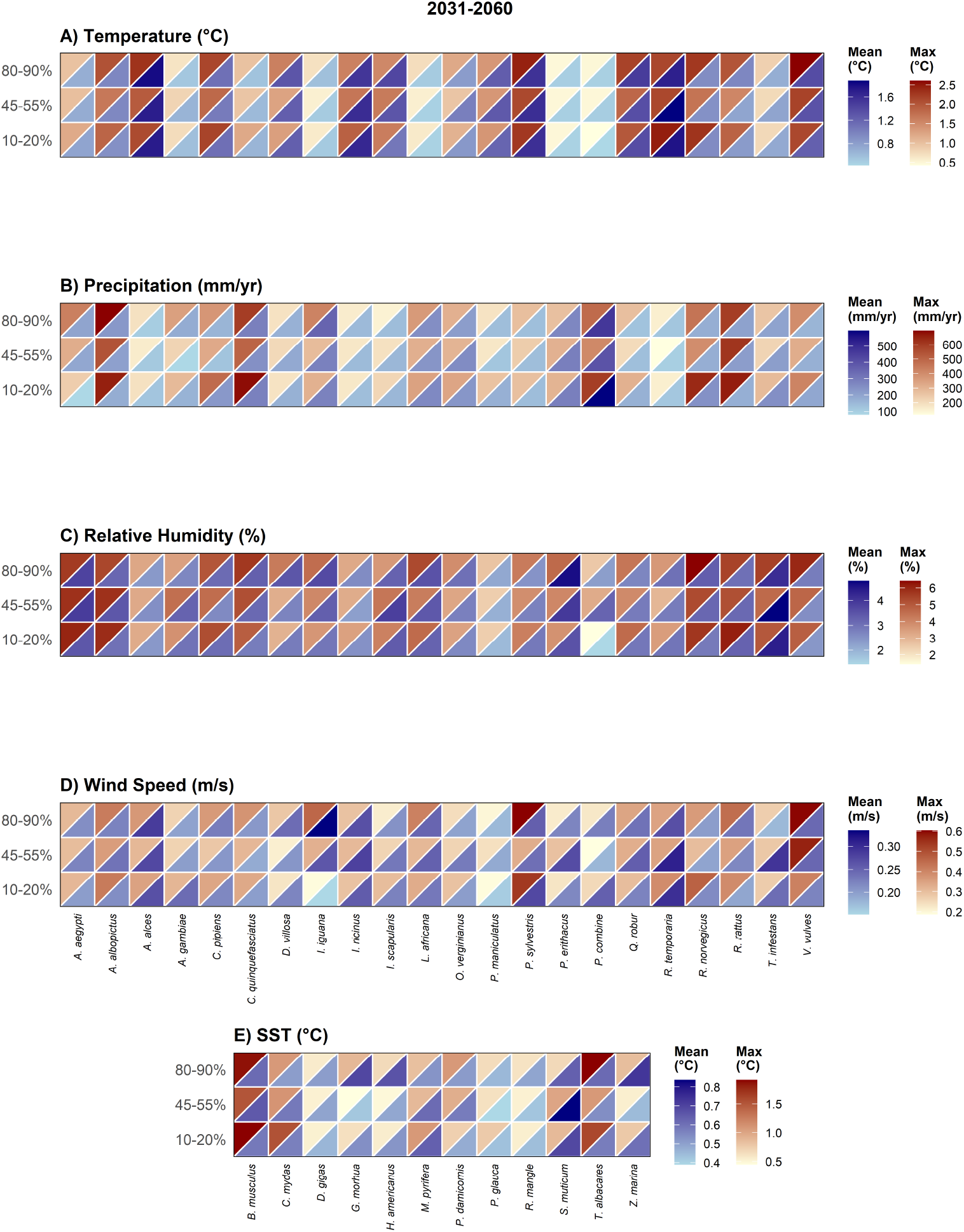
Internal-variability ranges across species and consensus bins for the period 2031–2060. Heat maps showing the relationship between species consensus and range of ICV of climate variables across initial-condition ensemble members for the 30-year annual average for 2031–2060. Across variables and species, internal-variability ranges overlap strongly among consensus bins, with no consistent ordering from low to mid to high consensus.

**Figure S6.**
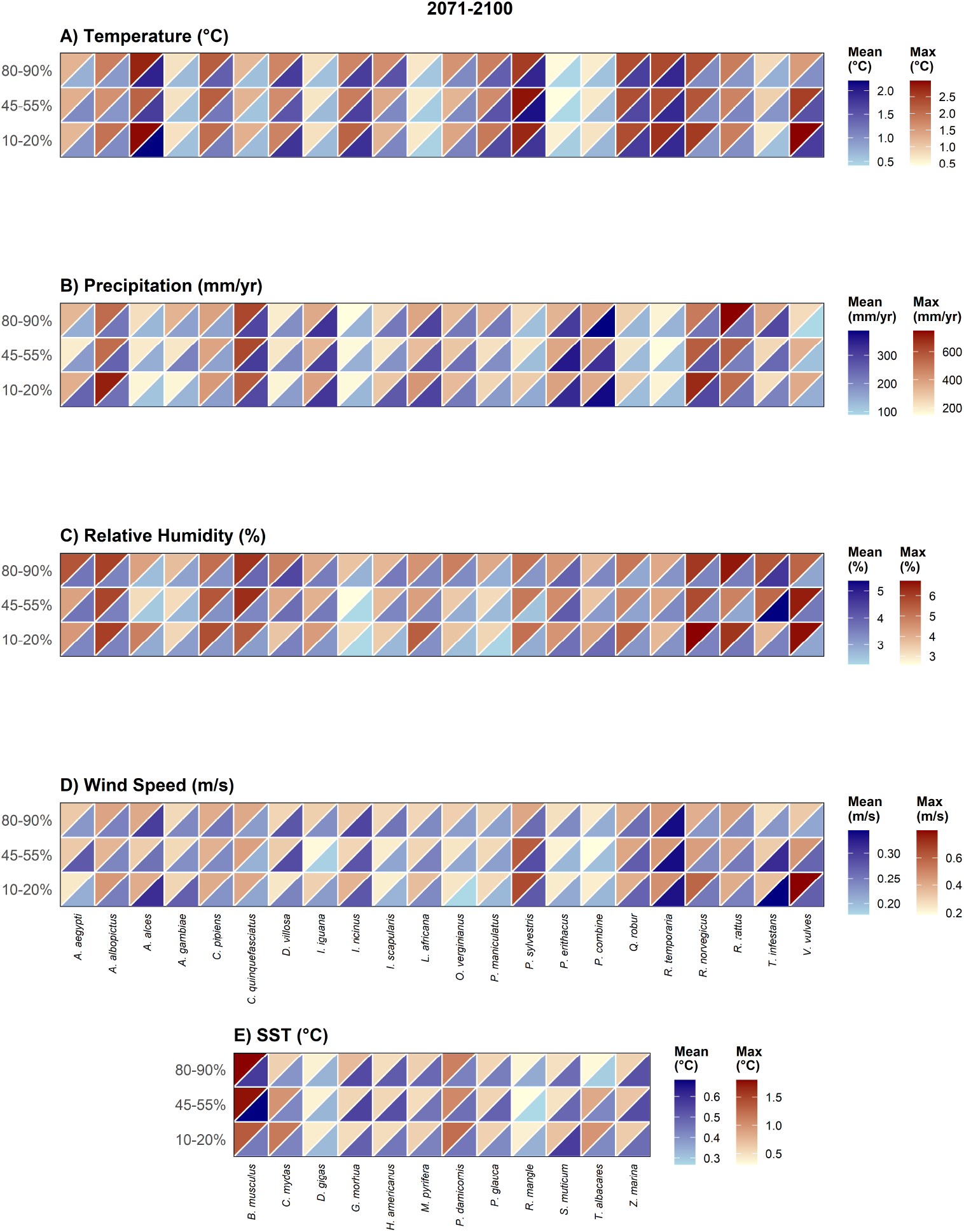
Internal-variability ranges across species and consensus bins for the period 2071–2100. Heat maps showing the relationship between species consensus and range of ICV of climate variables across initial-condition ensemble members for the 30-year annual average for 2071–2100. Across variables and species, internal-variability ranges overlap strongly among consensus bins, with no consistent ordering from low to mid to high consensus.

